# Use of phase plate cryo-EM reveals conformation diversity of therapeutic IgG with 50 kDa Fab fragment resolved below 6 Å

**DOI:** 10.1101/2023.12.21.571784

**Authors:** Hsin-Hung Lin, Chun-Hsiung Wang, Yi-Min Wu, Shih-Hsin Huang, Sam Song-Yao Lin, Takayuki Kato, Keiichi Namba, Naoki Hosogi, Chihong Song, Kazuyoshi Murata, Ching-Hsuan Yen, Tsui-Ling Hsu, Chi-Huey Wong, I-Ping Tu, Wei-Hau Chang

**Affiliations:** Institute of Chemistry, Academia Sinica, Taipei, Taiwan; Institute of Statistical Science, Academia Sinica, Taipei, Taiwan; Graduate School of Frontier Biosciences, Osaka University, 1-3 Yamadaoka, Suita, Osaka, Japan; JEOL Ltd., 1-2 Musashino 3-chome, Akishima, Tokyo, Japan; Exploratory Research Center on Life and Living Systems (ExCELLS) and National Institute for Physiological Sciences (NIPS), National Institutes of Natural Sciences, 38 Nishigonaka Myodaiji, Okazaki, Aichi, Japan; Genomic Research Center, Academia Sinica, Taipei, Taiwan; Institute of Physics, Academia Sinica, Taipei, Taiwan

**Author notes:** Institute of Preventive Medicine, National Defense Medical Center, New Taipei City, Taiwan (Shih-Hsin Huang). Cryo-EM Facility, College of Medicine, National Taiwan University, Taipei, Taiwan. Institute of Protein Research, Osaka University, Suita, Osaka, Japan (Takayuki Kato).

## Abstract

While cryogenic electron microscopy (cryo-EM) is fruitfully used for harvesting high-resolution structures of sizable macromolecules, its application to small or flexible proteins composed of domains like immunoglobulin (IgG) remain challenging. Here, we applied single particle cryo-EM to Rituximab, a therapeutic IgG mediating cancer cell toxicity, to explore its solution conformations. We found Rituximab molecules exhibited aggregates in cryo-EM specimens contrary to its solution behavior, and utilized a non-ionic detergent to successfully disperse them as isolated particles amenable to single particle analysis. As the detergent adversely reduced the protein-to-solvent contrast, we employed phase plate contrast to mitigate the impaired protein visibility. Assisted by phase plate imaging, we obtained a canonical three-arm IgG structure with other structures displaying variable arm densities co-existing in solution, affirming high flexibility of arm-connecting linkers. Furthermore, we showed phase plate imaging enables reliable structure determination of Fab to sub-nanometer resolution from ab initio, yielding a characteristic two-lobe structure that could be unambiguously docked with crystal structure. Our findings revealed conformation diversity of IgG and demonstrated phase plate was viable for cryo-EM analysis of small proteins without symmetry. This work helps extend cryo-EM boundaries, providing a valuable imaging and structural analysis framework for macromolecules with similar challenging features.

## Introduction

Antibodies, essential components of the immune system, are specialized immunoglobulin molecules designed to identify specific foreign substances, primarily proteins from pathogens such as viruses and bacteria. There exist five main classes of immunoglobulins: IgG, IgM, IgA, IgD, and IgE, with IgG being the most prevalent and having the smallest size. Upon binding to foreign molecular agents of a pathogen known as antigens, antibody molecules neutralize the invading pathogen. In the context of SARS-CoV-2 (COVID-19), individuals who have been infected can produce antibodies that specifically recognize the spike protein, preventing the virus from infecting additional cells. The specificity of these antibodies to SARS-CoV-2 proteins makes them valuable for testing, detection, and potential therapeutic use for seriously ill patients.

Therapeutic antibodies have been extensively developed for treating inflammation and cancer, and they are primarily in monoclonal form, for which traditional production relies on hybridoma technology [1] whereas recent trend for humanized variants involve screening from phage display libraries [2–4]. Notable therapeutic antibodies have been developed against Malignant B-cell lymphoma by targeting antigens derived from B-cell surface markers. One such marker is Cluster of Differentiation 20 (CD20), a membrane protein abundantly expressed on the surface of B cells throughout different stages towards maturation. Rituximab, an anti-CD20 monoclonal antibody of the IgG class, has been successfully employed for the molecular therapy of indolent B cell non-Hodgkin’s lymphoma (NHL) [5]. Rituximab binds to the CD20 receptor on malignant B cells, and mediates the recruiting of immune cells to the targeted tumor cell, thereby initiating the destruction of the cancer cell through secreted cytokines, a process commonly known as antibody-dependent cell-mediated cytotoxicity (ADCC) [6] (**Fig. 1**A). The intricate mechanism underlying this process has recently begun to unfold [7–9].

**Figure 1.**
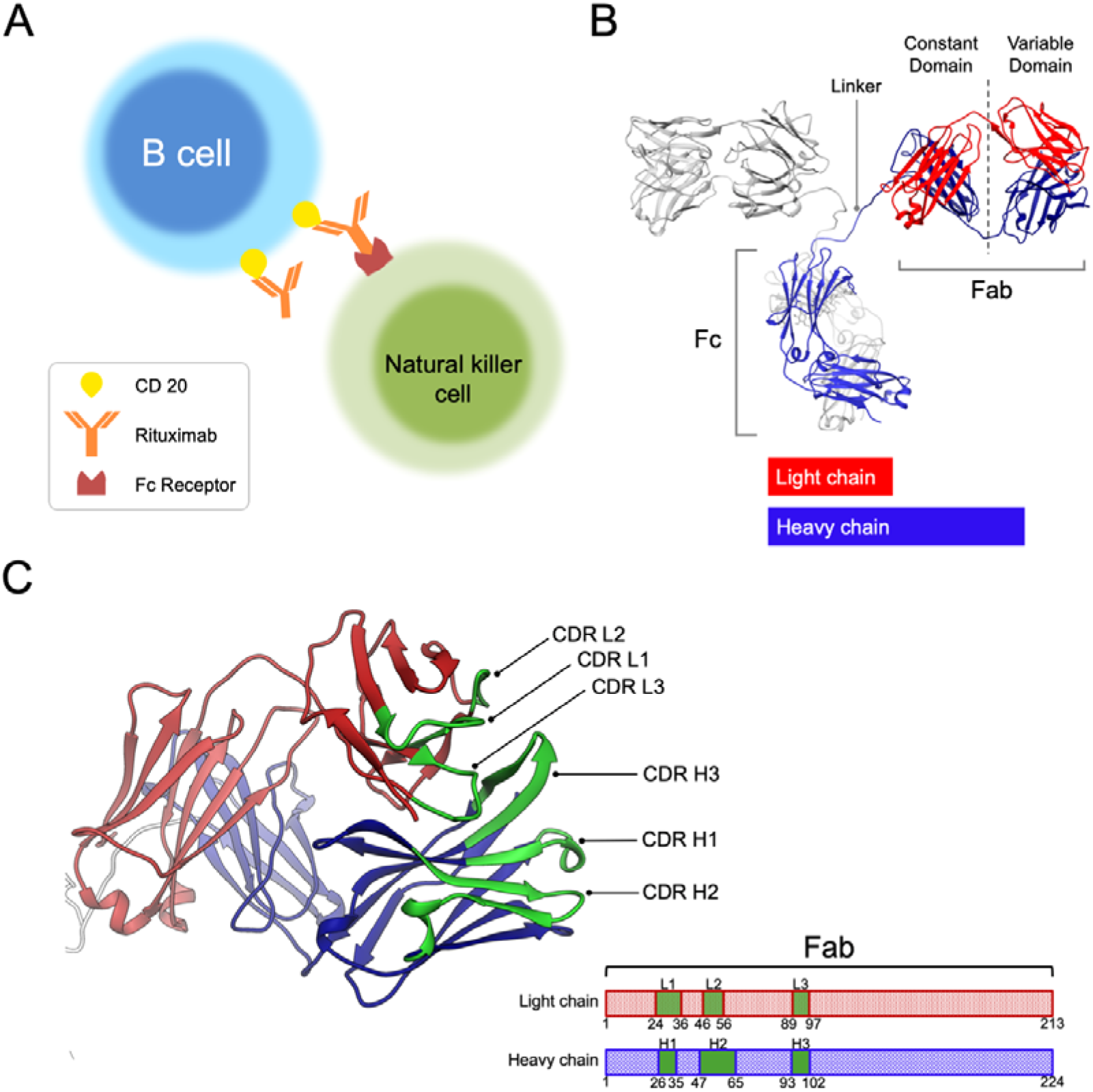
The action and structure of Rituximab. **(A)** Molecular basis of antibody-dependent cell-mediated cytotoxicity against malignant B cells. Rituximab binds to CD20 receptors on malignant B cells via its Fab and recruits Nature Killer cells via its Fc. Cytokines released (not shown) by Nature Killer cell can affect the nearby B cell to induce its apoptosis. **(B)** An atomic model of IgG (PDB: 1IGT). **(C)(D)** The structure (PDB: 4KAQ) and sequences of Rituximab heavy chain light chain in Fab—highlighted are the sequences of the complementary determining region (CDR) for recognizing the epitope in CD20.

The dual ability of IgG to target tumor cells and recruit immune cells is deeply rooted in its exceptional structure. Through X-ray crystal analysis [10], an intact IgG molecule reveals a Y-shaped architecture built from four polypeptide chains (**Fig. 1**B). This structure comprises two identical Fab domains with Fab representing “antigen-binding fragments”, and one Fc domain with Fc denoting “crystallizable fragment”. Each Fab domain possesses an antigen-binding site, composed of six complementarity-determining regions (CDRs) (**Fig. 1**C) in the variable region (Fv), responsible for recognizing antigens, e.g. receptor proteins on tumor cells. The Fc region can bind to various receptor molecules on immune cells, conferring effector functions. The variable region (Fv) of Fab consists of a pair of variable domains, VH and VL, forming part of the heavy and light chains linked by disulfide bonds (**Fig. 1**B).

While the snapshot structure of IgG has been known (**Fig. 1**B), potential dynamics of IgG have been inferred based on large thermal factors observed for the Fab-Fc linking region in X-ray diffraction analysis. The focus of structural analyses of antibodies has therefore gradually shifted towards assessing the conformational flexibility in connection with more sophisticated functional aspects. In particular, there is a growing pharmaceutical interest in understanding the conformational mechanisms that enhance the therapeutic efficacy of antibodies [11]. This question could be better addressed using techniques that do not constrain antibody conformations in crystals. Compared to X-ray crystallography, cryo-electron microscopy (cryo-EM) is virtually a solution technique capable of accessing structural polymorphism in solution because the recorded images represent a body of mixed data of all co-existing solution conformations [12]. In addition, cryo-EM imaging method is versatile in that it accommodates nearly all sample conditions in response to physicochemical or biochemical manipulations. Cryo-EM with tomographic reconstruction has been previously utilized to reveal structural variations of IgG [13], albeit at low resolutions due to technological limitations.

The recent advent of direct electron detectors [14–15] and Bayesian image analysis algorithms [see 16] have together transformed single-particle cryo-EM into a high-resolution method for determining the structures of sizable and rigid biological macromolecules (>100 kDa) without the need of crystals, particularly beneficial for large protein complexes. Notably, various IgG-receptor complexes [17–19], including IgG complexed with CD20 [8–9], have been determined at medium to high resolution using the single-particle approach. It is important to note that the success of these celebrated cryo-EM studies has enjoyed the signals provided by the large mass from the part of antibody-binding partners. Even in the case of artificially designed Legobody [20], the antibody is decorated by nanobody and maltose, together exceeding a molecular weight of 100 kDa. The current mass limit of high-resolution single particle cryo-EM is held by a 60 kDa protein with symmetry [21].

In spite of carrying molecular weight of 150 kDa above the mass limit threshold, IgG molecule is not conducive to achieving high-resolution structures with use of single-particle approach because it is a flexible protein composed of small domains (50 kDa) that appear as obscure dots in vitreous ice. Traditionally, visual identification of IgG by electron microscopy has utilized negative-stain chemicals for contrast enhancement [22–23], which may incur structural artifacts due to dehydration. Thus, to clearly visualize dots of IgG may invoke use of advanced phase contrast techniques compatible with cryo-EM. [24,25].

In this study, we explored use of phase plate to perform cryo-EM imaging of Rituximab to explore its conformational dynamics. During our cryo-EM imaging experiments, we surprisingly encountered the issue of Rituximab aggregation. Utilizing a non-ionic detergent (DDM), we rendered Rituximab molecules mono-dispersed, and adopted phase plate imaging to amend the cryo-EM contrast worsened by DDM, leading to the first cryo-EM reconstruction of therapeutic IgG in isolation. This reconstruction showcases a canonical three-arm IgG structure, with co-existing structures lacking one or two arms, indicating an extremely high degree of flexibility associated with the arm-connecting linker. Furthermore, we demonstrated that the use of phase plate enabled single-particle structural determination of Fab fragment of 50 kDa to below 6 Å from ab initio.

## Results

### Aggregation of Rituximab molecules found in cryo-EM specimen

Maintaining good solubility for therapeutic IgG at high concentrations is crucial for the safe use of this protein drug. To achieve mono-dispersity for therapeutic IgG, various undisclosed additives are incorporated into its formulation by pharmaceutical companies [see 26]. To evaluate the solution behavior of a commercially supplied Rituximab that is biochemically pure (**Fig. 2**A), we employed dynamic light scattering (DLS). The DLS measurements showed the vendor-supplied Rituximab molecules (**Fig. 2**B), whether undiluted or diluted with phosphate-buffered saline (PBS) or de-ionized water, predominantly exhibited a sharply defined size-distribution centered around 7 nanometers (**Fig. 2**C & D), affirming that the Rituximab molecules were largely homogeneous in solution.

**Figure 2.**
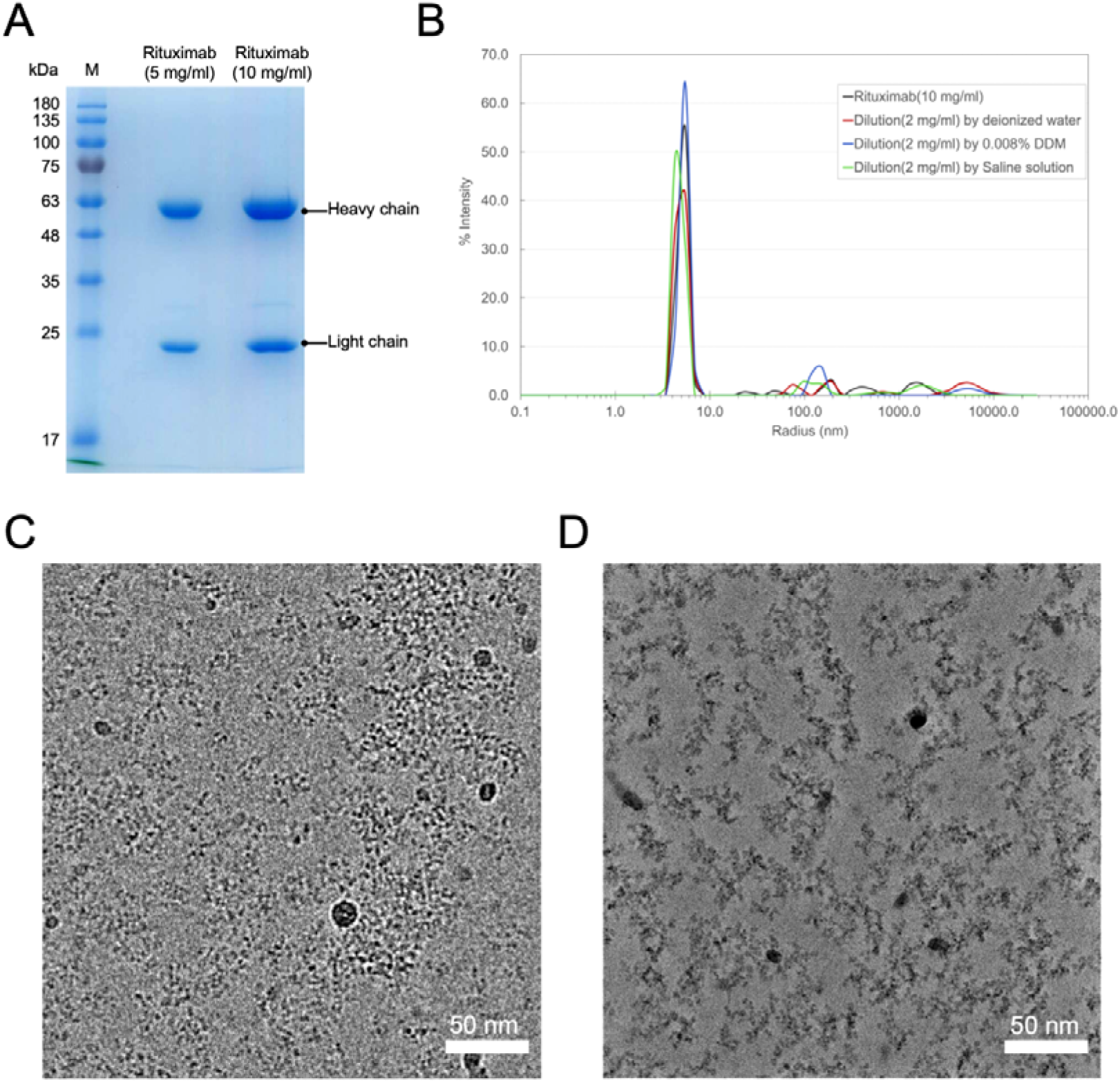
Biochemical and biophysical characterization of Rituximab. **(A)** SDS PAGE gel of Rituximab loaded with two concentrations (10 mg/ml & 5 mg/ml), showing the heavy and light chains. **(B)** Dynamical light scattering of Rituximab, undiluted (10 mg/ml) (black), diluted to 2 mg/ml, with deionized water (red), DDM (0.008%) (blue), and saline (green) respectively. The Y axis represents scattering intensity and X-axis molecular size. **(C) & (D)** Cryo-EM images of Rituximab (diluted to 2 mg/ml only with deionized water) recorded without and with phase plate. (C) is by conventional defocusing imaging (by Talos Arctica, Thermo Fisher Scientific, USA), and (D) is with phase plate (hole-free phase plate on F-200, JEOL, Japan).

However, as we performed cryo-EM imaging of specimens made from this vendor-supplied Rituximab via blotting and plunge-freezing, we repeatedly observed Rituximab exhibited aggregates, contrary to our expectations based on the DLS measurements. Such unexpected behavior of this IgG in thin liquid layer severely hindered our structural investigation using single-particle method.

### Zernike phase plate facilitates visualization of DDM-embedded Rituximab

Speculating that the aggregation might be associated with the air-water interface issues [27–28], we introduced a non-ionic detergent into the Rituximab-diluting buffer since this approach had effectively mitigated the air-water interface issues [29]. We chose n-Dodecyl-beta-Maltoside (DDM) as it had been proven compatible with Rituximab, given its previous use in cryo-EM studies of Rituximab in complex with CD20 [8,9]. Yet, the presence of DDM was found to significantly impair cryo-EM image quality. In the past, Zernike phase plate (ZPP) imaging was successfully adopted to visualize DDM-solubilized membrane proteins [30]. We therefore resorted to use of ZPP to facilitate cryo-EM imaging of Rituximab in DDM. Fig. 3A displays a number of ZPP micrographs (see **Table 1**) that clearly show sparse Rituximab molecules (0.5 mg/ml) as Y-shape particles (red circles in **Fig. 3**A) in thick DDM background (**Fig. S2**) without signal enhancement using 2D classification. Further 2D classification enabled automated sorting of various orientations of Rituximab molecules (**Fig. 3**B). In our control study that employed conventional cryo-EM, we had encountered much greater difficulties in visualizing Rituximab in DDM (**Fig. S1**). Those findings indicated that cryo-EM studies involving use of contrast impairing additive (**Fig. S2**) can benefit from phase plate imaging.

**Figure 3.**
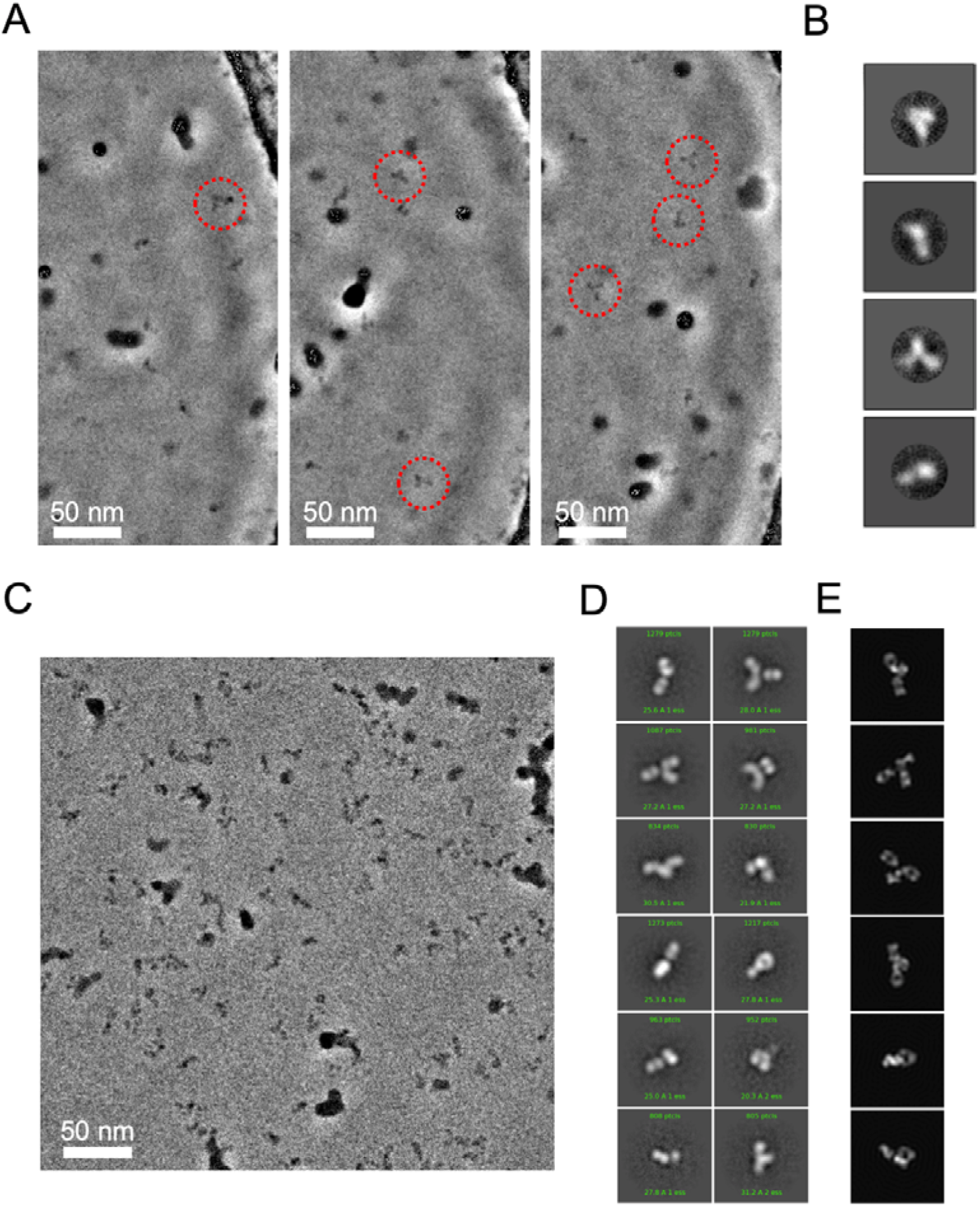
Phase plate cryo-EM imaging of DDM-solubilized Rituximab. **(A)** ZPP images show Rituximab (∼0.5 mg/ml with 0.008% DDM) molecules are sparsely distributed as denoted in red circles. **(B)** Four representative 2D class averages generated from picked ZPP Rituximab particle images. **(C)** VPP images of Rituximab (∼2.0 mg/ml with 0.008% DDM). **(D)** 2D class averages generated from picked VPP Rituximab particle images. **(E)** Representative 2D re-projections from an IgG crystal structure (PDB: 1IGT) for comparison.

**Table 1.**
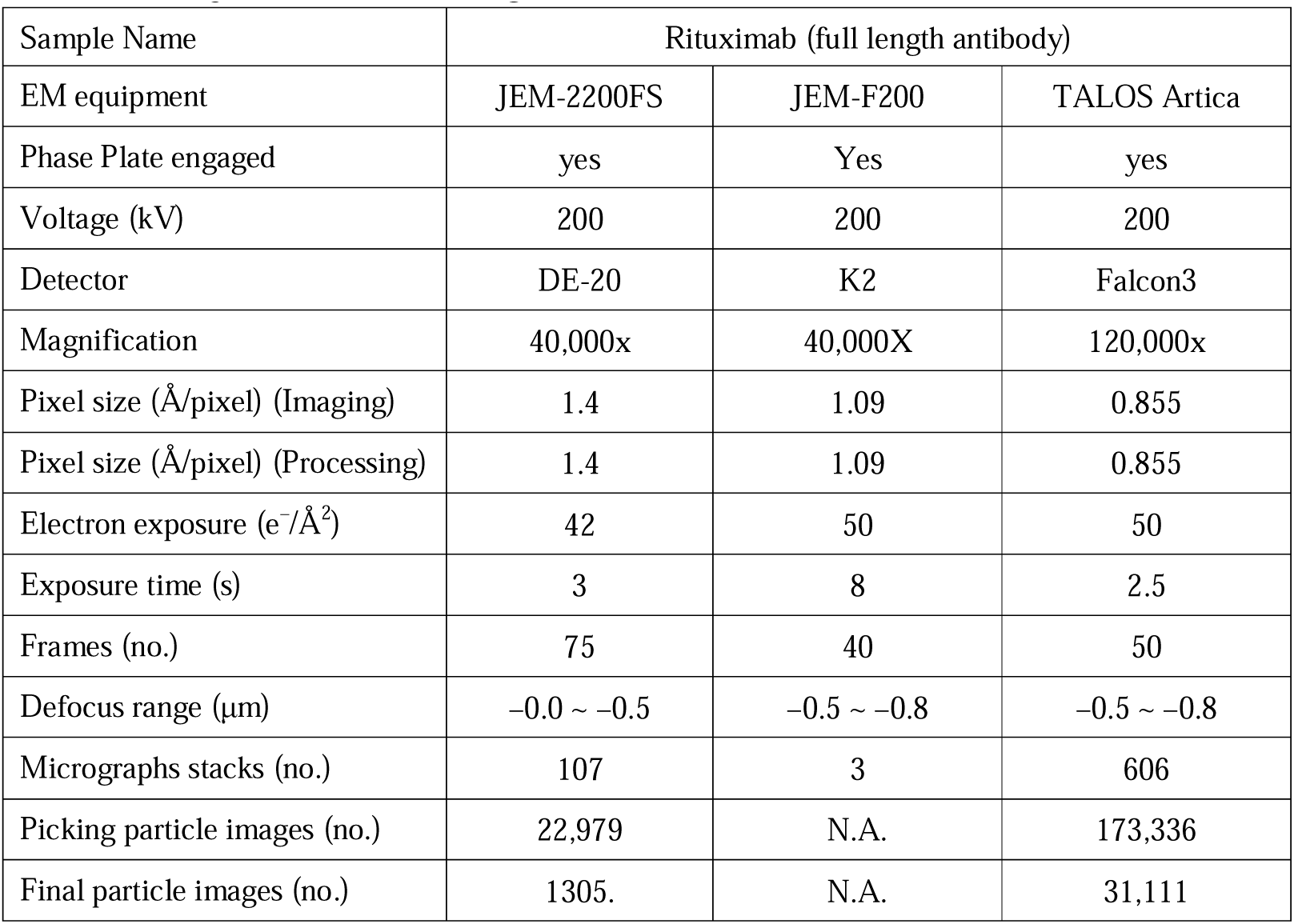
Cryo-EM used for IgG studies.

| Sample Name | Rituximab (full length antibody) |  |  |
| --- | --- | --- | --- |
| EM equipment | JEM-2200FS | JEM-F200 | TALOS Artica |
| Phase Plate engaged | yes | Yes | yes |
| Voltage (kV) | 200 | 200 | 200 |
| Detector | DE-20 | K2 | Falcon3 |
| Magnification | 40,000x | 40,000X | 120,000x |
| Pixel size (Å/pixel) (Imaging) | 1.4 | 1.09 | 0.855 |
| Pixel size (Å/pixel) (Processing) | 1.4 | 1.09 | 0.855 |
| Electron exposure (e<sup>-</sup>/Å<sup>2</sup>) | 42 | 50 | 50 |
| Exposure time (s) | 3 | 8 | 2.5 |
| Frames (no.) | 75 | 40 | 50 |
| Defocus range (μm) | -0.0 ~ -0.5 | -0.5 ~ -0.8 | -0.5 ~ -0.8 |
| Micrographs stacks (no.) | 107 | 3 | 606 |
| Picking particle images (no.) | 22,979 | N.A. | 173,336 |
| Final particle images (no.) | 1305. | N.A. | 31,111 |

### Volta phase plate imaging reveals fine structures in Rituximab arms

While ZPP offers drastic contrast due to full restoration of low-resolution spatial information (**Fig. S3**), its usage with near in-focus conditions without CTF correction restricts the resolution to nanometer [30]. In keeping our aim in pursuit of structural analysis of Rituximab to higher resolution using PP imaging, we turned to the Volta phase plate (VPP) [31–33]. Compared to ZPP as pre-fabricated micron-size pinholes on a thin carbon film (24 nm for 200 kV), VPP is a hole-free solution where it is in situ made on a thinner continuous carbon film (12 nm) by exposure to irradiating electrons. Importantly, use of VPP with slight defocus allowed for CTF correction. This approach has been shown to achieve near-atomic resolution with benchmark specimens with high symmetry [34]. Practically, use of VPP is facilitated by its availability on modern cryo-EM with automation that can easily scale up the number of images required for analyzing structures with low or no symmetry, and also for investigating the relationship between resolution and number of particles [35].

Guided by the sample conditions learned from our aforementioned pilot ZPP studies, we optimally diluted Rituximab with a buffer containing DDM (0.008%) to 2 mg/ml, and collected 606 movies from one specimen using VPP on a modern cryo-EM (**Table 1**) in an overnight session. As shown in a VPP micrograph (**Fig. 3**C), the feature of Y-shaped structures of Rituximab IgG could be recapitulated. Remarkably, further 2D class classification of the CTF-corrected VPP images disclosed fine details within the arms (**Fig. 3**D), and gave average images consistent with projections from IgG crystal structure (PDB: 1IGT) (**Fig. 3**E).

### 3D reconstruction of Rituximab from VPP images

To derive the 3D structure of Rituximab from VPP images, we performed single-particle analysis with two widely used algorithms, RELION [16] and cryoSPARC [36]. Initially, we utilized the Blob Picker program with a particle diameter set at 150 Å (15 nm) for automatic particle picking from the 606 micrographs that were motion-corrected, dose-weighted, and CTF-corrected. Those picks then underwent one round of reference-free 2D classification using RELION [16] to yield classes exhibiting clear class average images, which we used as templates for Template Picker on RELION to re-perform particle picking and obtained a total of 173,336 particles. Multiple rounds of reference-free 2D classification were subsequently performed to yield a presumably homogeneous subset of 31,111 particles (**Table 1**).

While X-ray structure of IgG could be adopted as a reference model to facilitate 3D reconstruction, we took an ab initio approach for those 31,111 particles using ab initio model generation on cryoSPARC (v4.1) [36]. Doing so was meant to avoid potential “Einstein from noise” [37] from use of external model. Notably, different tosses with “ab initio model generation” gave distinctly different results. As shown in **Fig. S4**, a low-resolution model with full density of three IgG arms could be occasionally obtained. Using this three-arm model as the reference, we conducted global 3D refinement to optimize the angle and translation alignment parameters of each IgG image in the set of 31,111 particles to obtain a canonical three-arm structure of Rituximab in solution (**Fig. 4**A). As shown in **Fig. 4**B, the angle distribution is largely uniform with reasonable coverage. This Rituximab structure exhibits overall resolution of 8.45 Å based on the gold-standard FSC at a 0.143 cutoff (**Fig. 4**C) where local resolution analysis [38] showed the resolutions varied in the range of 8.3 to 17.3 Å (**Fig. 4**D). One of the arms could be recognized as Fab as it displayed a characteristic Fab structure with two lobes still connected.. Notable features of this IgG structure include: (i) the dyad axis between two Fab domains does not coincide with that of Fc, which is slightly tilted. (ii) the Fab-Fc connecting linker in this canonical form is rigid; (iii) the resolutions within Fab or Fc domain are not uniform, indicative of internal flexibility. Interesting, as we performed rigid-body docking of crystal structure (PDB: 1IGT) into this Rituximab cryo-EM reconstruction, we found it significantly deviated from the crystal structure of an IgG in the same sub-class To further elucidate the possible movements that cause such discrepancy, we docked individual crystal structures of Fab and Fc into the corresponding region in our cryo-EM map as separate entities (**Fig. 4**E), and compared the the fitted Fc with that in the X-ray structure of an IgG (PDB: 1IGT). We surprisingly discovered pronounced rotation and rocking movements with the Fc domain (**Fig. 4**F).

**Figure 4.**
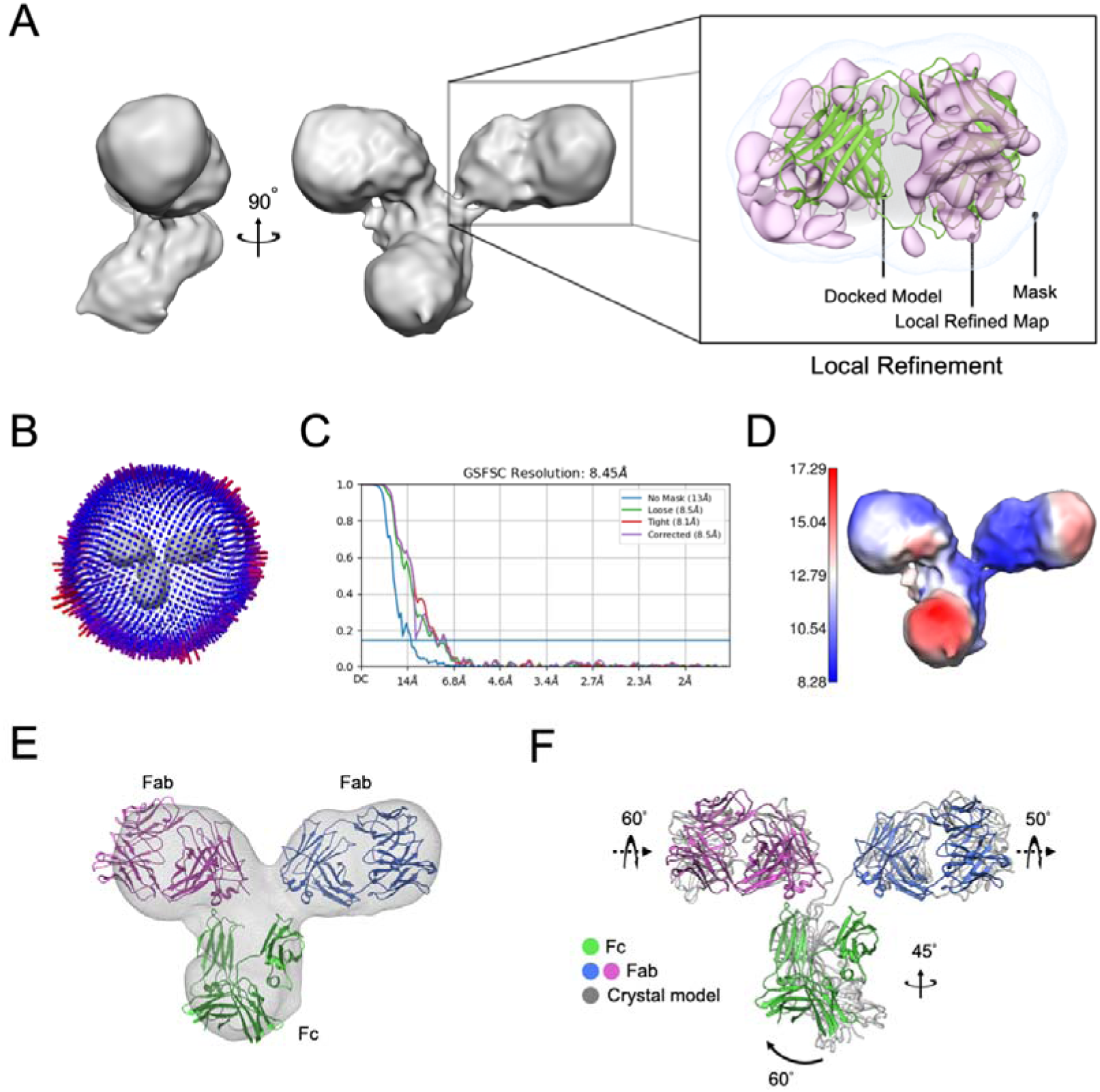
3D reconstruction of Rituximab from Volta phase plate (VPP) cryo-EM images. **(A)** 3D cryo-EM reconstruction of Rituximab with Fab density further focused refined as shown in the box. **(B)(C)(D)** angular distribution of IgG particle orientations, FSC from cryoSparc for resolution estimation, and local resolution maps. **(E)** Docking of Fab and Fc as individual domains into our cryo-EM reconstruction. **(F)** Comparison of the pseudo atomic model generated from (E) to that of a crystal structure (PDB: 1IGT).

We then sought to improve the resolution of Fab arm by using focused refinement. This effort seemed to completely resolve Fab into two separate lobes (shown in the box of **Fig. 4**A), but the resulting spurious densities had tainted the map reliability (**Fig. 5**A). Of note, the same effort applied to Fc gave little improvement (data not shown).

**Figure 5.**
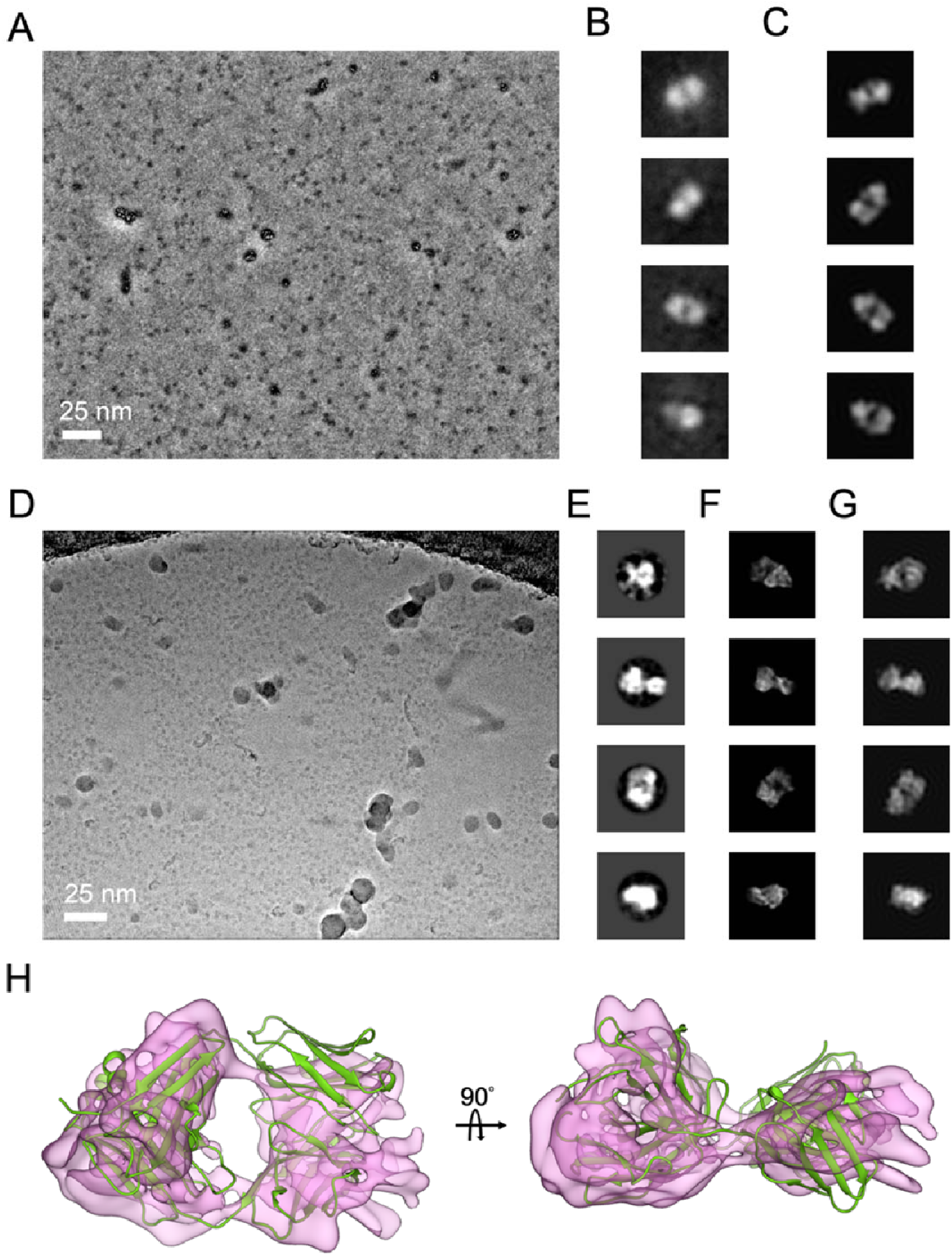
Phase plate imaging of Fab fragment. **(A)** ZPP Fab particles seen as dense “black” dots. **(B),(C)** Five representative 2D class averages and low-pass re-projections of crystal structure (PDB:4KAQ) in similar views. **(D)** VPP Fab particles seen as “gray” dots. **(E),(F),(G)** Five representative 2D class averages, re-projections from 4KAQ, and cryo-EM structure from (H). **(H)** Cryo-EM reconstruction of Fab fragment with docking of 4KAQ.

To further explore the conformational diversity arising from linker mobility, we further performed 3D variation analysis (3DVA) [39] on this 31,111 particle set. Remarkably, the 3DVA analysis further categorized Rituximab particles into four 3D classes: Class_1 (9.26%) is with only one Fab arm, Class_2 (13.74%) and Class_3 (20.35%) are with three arms, and Class_4 (56.65%) is with two arms where one Fab seems missing (**Fig. S4**). These results show that Rituximab molecules exhibit highly diverse conformations.

### Cryo-EM structural determination of Fab fragment

To better resolve Fab, we explored the possibility of analyzing it as an isolated object. To this end, we derived Fab fragments from Rituximab by enzymatic cleavage [40]. It is noted that Fab domain has a molecular weight of 50 kDa, almost falling outside the range suitable for high-resolution single-particle cryo-EM analysis. Of note, haemoglobin (64 kDa, C2 molecule) was analyzed using conventional cryo-EM to 3 Å resolution while the same approach to protein kinase A (PKA) without symmetry (43 kDa, C1 moleucle) only led to 6 Å resolution [21].

In this single particle pursuit of Fab, we compared phase plate to conventional cryo-EM. Recognizing the benefit of ZPP for visualizing small particles [25,41–42], we initially employed ZPP cryo-EM to capture images of the Fab fragment on a multi-purposed cryo-EM (**Table 2**). Shown in **Fig. 5**A is ZPP cryo-EM micrograph that shows countless small “dots” representing particles with size expected from Fab. Remarkably, reference-free 2D classification from those “dots” revealed the characteristic two-lobe structure of Fab (**Fig. 5**B). We were convinced that these dots were indeed Fab molecules as those class averages are in agreement with views projected from known Fab structure (PDB: 4KAQ) that were low-pass filtered (**Fig. 5**C),. This line of study demonstrates that ZPP contrast can help detecting small proteins in highly noise associated with low-dosed cryo-EM, and enable accurate alignment of the “featureless” particle images, providing an experimental proof with sub-100 kDa protein to support the ZPP benefits predicted by modeling studies [41–42] prior to the cryo-EM revolution,

**Table 2.** Cryo-EM used for Fab studies.

| Sample Name | Fab |  |  |
| --- | --- | --- | --- |
| EM equipment | JEM-2200FS | Titan Krios |  |
| Phase Plate engaged | Yes | Yes | no |
| Voltage (kV) | 200 | 300 |  |
| Detector | DE-20 | K3 |  |
| Magnification | 40,000x | 105,000x |  |
| Pixel size (Å/pixel) (Imaging) | 1.4 | 0.415 |  |
| Pixel size (Å/pixel) (Processing) | 1.4 | 0.83 |  |
| Electron exposure (e<sup>-</sup>/Å<sup>2</sup>) | 42 | 50.56 |  |
| Exposure time (s) | 3 | 1.62 |  |
| Frames (no.) | 75 | 50 |  |
| Defocus range (μm) | -0.0 ~ -0.5 | -0.5 ~ -0.8 | -1.6 ~ -3.3 |
| Micrographs stacks (no.) | 55 | 956 | 10,166 |
| Picking particle images (no.) | 44,967 | 288,931 | 12,650,536 |
| Final particle images (no.) | 7,080 | 72,035 | 305,246 |

To scale up Fab particle images for 3D reconstruction, we turned to use VPP on our 300 kV instrument to enjoy automated data collection with a direct electron camera operated at sub-0.5 Å pixel resolution. We collected 950 VPP cryo-EM movies in a one-day session. In a VPP image shown in **Fig. 5**D, one can observe numerous small “dots” with a size consistent with that of Fab, albeit the contrast appears to be weaker than that achieved with ZPP. Through several rounds of reference-free 2D classification, we selected approximately 70,000 “good” VPP particles from the total of approximately 0.28 million picked particles that were CTF-corrected (**Table 2**). Compared to the ZPP 2D class averages, the VPP 2D class averages gave more structural details (**Fig. 5**E), corroborated by comparison to corresponding projection views from the crystal structure (PDB: 4KAQ) that were without low-pass filtering (**Fig. 5**F). We sought to determine whether or not a 3D structure of Fab could be generated from ab initio using these ∼70,000 selected particles. By using cryoSPARC (v4.1)[36], we found that approximately one out of three tosses could successfully hit the target—a low-resolution structure with two lobes, which we then adopted as a reference model for further 3D refinement. 3D refinement using VPP images led to a typical tertiary structure of Fab (**Fig. 5**G & **6**B), arriving at approximately 6 Å resolution (**Fig. 6**E). While the secondary structure of the β-sheet in this Fab reconstruction could not be resolved at this resolution, the reliability of this Fab map is confirmed by rigid-body docking of known Fab X-ray structure (PDB: 4KAQ) (**Fig. 5**G).

**Figure 6.**
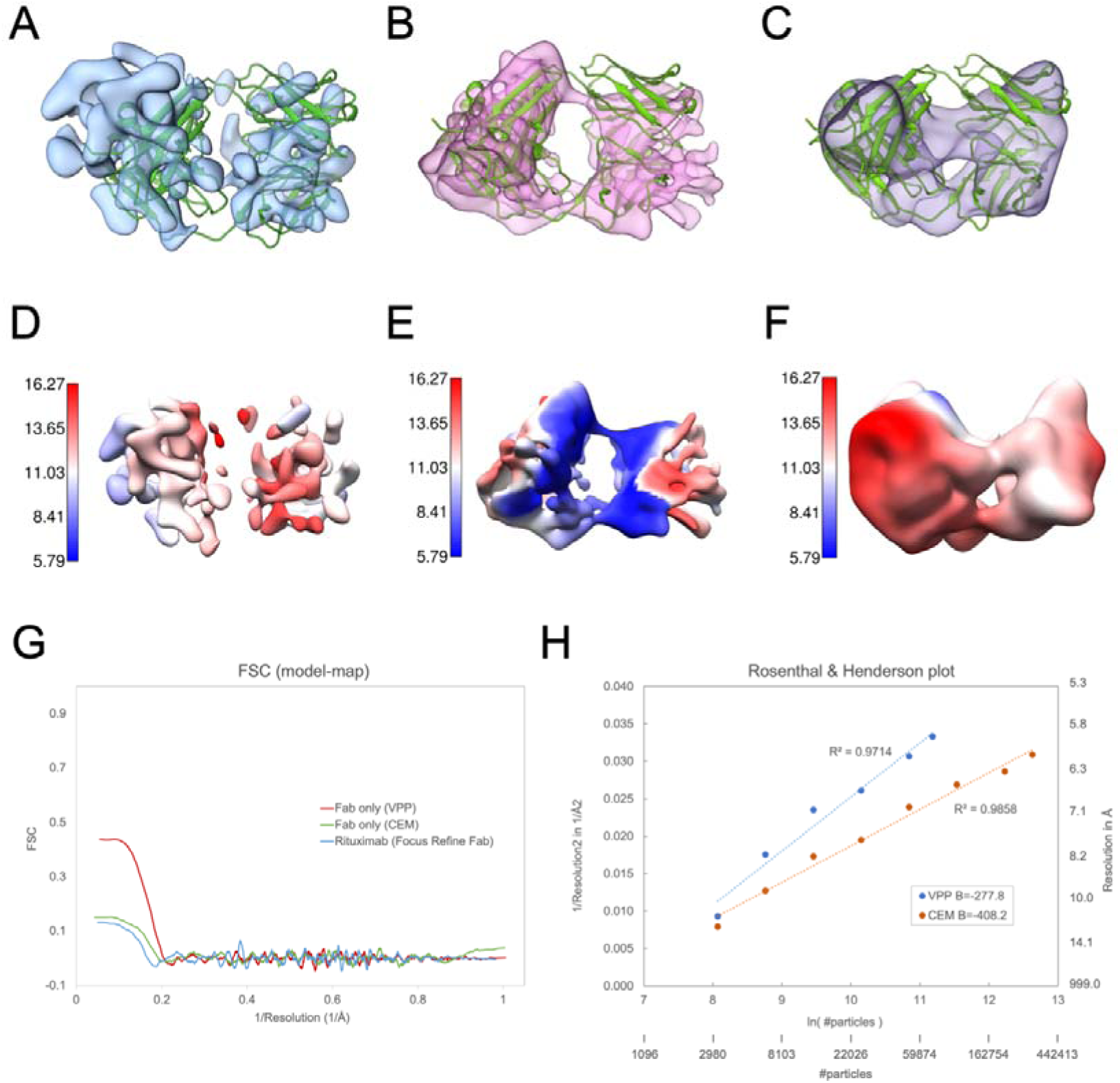
Comparison of three Fab reconstructions. **(A)** Fab from focused refinement of its density in VPP IgG images. (**B)** VPP Fab. **(C) CEM** Fab. **(D),(E),(F)** local resolution maps of (A)(B)(C). **(G)** Map-to-model FSC for (A)(B)(C). **(H)** A Rosenthal & Henderson (R & H) plot describing the resolution versus number of particles, VPP (blue) and CEM (red). Our R & H analysis was performed with cyoSPARC based on the sampling scheme described by the script “bfactor_plot.py” in Relion. Uncovering that in our case the degree of filtering for an initial model strongly affected the reported resolution of final reconstruction when less than 3,000 particles were used, we started with 3,200 particles as the smallest randomly sampled subset and doubled the particles in the analysis sequence until all particles were exhausted.

To further compare conventional defocusing imaging (CEM) with PP imaging, we collected 10,166 micrographs in a 48h session to yield a very large CEM dataset containing a total of ∼12 million particles (**Table 2**). Through extensive reference-free 2D classification (**Fig. S**5), we selected 300,000 ‘good” particles for 3D reconstruction from ab initio. This effort led to a CEM structure with two lobes (**Fig. 6**C & F) in spite of all difficulties where this structure is with lower quality than that calculated from VPP images (**Fig. 6**B & E). We noticed the CEM particle yield from use of reference-free 2D classification was only 2.5%, which was much lower than the figure for VPP images (25%).

We then studied how resolution might improve as particle number increased with Rosenthal & Henderson analysis while using cryoSPARC for particle sampling and 3D reconstruction. For the case of CEM particle images, this analysis reveals that a fairly large B-factor measured around 400 Å^2^ (**Fig. 6**H). This value contrasts to 100 Å^2^ [34] or smaller typically observed for large or rigid cryo-EM structures determined to near atomic resolution or better, and reflects the fact that CEM images of small proteins are highly prone to mis-alignment with current algorithms [21]. In comparison, the B-factor for VPP images was reduced to 270 Å^2^ (**Fig. 6**H), demonstrating significant improvement in the accuracy of image alignment with increased phase contrast [41–42]. In addition, this comparison of VPP and CEM (**Fig. 6**G) shows that using CEM to target similar resolution reached by VPP, many more particles are required, affirming another benefit of using PP imaging for single particle structure determination of small protein particles, particularly those without symmetry [41].

## Discussion and conclusions

Here, we report the first cryo-EM structure of an immunoglobulin G (IgG) in isolation with sub-nanometer resolution. IgG is recognized for its distinctive three-blob or Y-shape architecture, comprising four polypeptide chains whose atomic details have long been elucidated through X-ray crystal diffraction analysis where vast diversity of IgG conformations in solution primarily arises from the movement of the long Fab-Fc connecting linkers. Understanding IgG conformations is crucial as it underscores the structural basis of its function. Tuning IgG conformations through molecular engineering has become an active area in the development of therapeutic IgG. However, an effective and efficient approach to characterizing solution IgG conformations in detail is on demand. Along this line, our single-particle work here has made further advance with improved resolution from previous work of Sandin et al that dissected IgG structural variations using cryo-electron tomography (cryo-ET) [13] with an innovative likelihood-based de-noising method to segment individual IgG densities from extremely noisy tomograms..

Single-particle cryo-EM has become a high-resolution method for studying many macromolecules without the need for crystallization. Compared to cryo-ET, single-particle approach is an averaging technique that it is less sensitive to structural variation. Nonetheless, companion structural variation analysis allows capturing structural polymorphism in hydrated state [12,39], even at medium resolutions. Given that IgG is a flexible protein composed of small domains (50 kDa) and thus considered as an unfavorable target for high-resolution cryo-EM analysis, we aimed to apply single-particle cryo-EM to Rituximab, a therapeutic IgG, to aim for medium-resolution information for exploring its solution conformations.

We encountered significant challenges at the specimen level—Rituximab persistently exhibited aggregation in the cryo-EM sample, and overcame this aggregation issue by employing a non-ionic detergent, successfully dispersing it into isolated particles and making it amenable to a single-particle approach. The use of detergent led to a severe reduction in protein-to-solvent contrast, significantly compromising image quality, for which we adopted PP imaging to rectify the impaired contrast. This unprecedented contrast allows direct identification of individual IgG molecules in raw images on-the-fly without the need of further enhancing signal-to-noise ratio (SNR) via post-imaging 2D image averaging.

During the structural analysis of approximately 30,000 IgG VPP particles, we encountered challenges related to structural heterogeneity in the dataset. Further 3D structural variation analysis revealed the presence of several co-existing structures, some with three arms, and some others lacking one or two arms, reflecting the flexibility of the linkers and the dynamics of Rituximab IgG in solution Notably, although three-arm IgG structures were previously observed using negative-stain electron microscopy [22–23], those results were not free from conformation distortion induced by dehydration.

Given that one of the central goals of structural research on IgG is to gain details of the Fab [43], particularly those in the CDR regions to understand how it recognizes epitopes, we employed focused refinement to enhance the resolution of Fab domain in IgG. While this effort seemed to resolve Fab from one connecting bulk into two separate lobes, the resulting spurious densities had suggested the map could have been over-fitted (**Fig. 6**A). To overcome these limitations, we pursued resolution for Fab (50 kDa) as an isolated entity. Considering that 50 kDa is near the lower mass limit for high-resolution single cryo-EM methods, as demonstrated by Herzik Jr et al [21] in the seminal work targeting sub-100 kDa proteins using a conventional defocusing approach, the achievable resolution of Fab remains uncertain, and the necessity of using phase plate imaging remains to be determined.

Our single particle cryo-EM study utilizing the Volta phase plate (VPP) to investigate Fab cleaved from Rituximab IgG showed that with a moderately sized particle set of approximately 70,000 good particles selected from a total of 950 micrographs a Fab reconstruction with resolution slightly below 6 Å could be achieved from ab initio. Although this map still fell short of resolving the β-strands within, it showcased characteristic Fab lobes and was more reliable than that obtained from focused refinement. Our attempts to dock the Fab X-ray crystal structure (4KAQ) into this Fab cryo-EM reconstruction supported the reliability of this structure. We noticed that in the case of protein kinase A (PKA) [21] determined to similar resolution by single particle CEM, the inner features representing secondary structures were more easily interpretable as PKA was primarily composed of α-helices. In comparison, our study of Fab using conventional defocusing cryo-EM (CEM) without PP showed CEM required a much larger dataset (300,000 good particles selected from ∼12 million particles across ∼10,000 micrographs) to achieve a reconstruction for Fab while the map quality was lower (**Fig. 6**F), as also evidenced by the map-to-model FSC (**Fig. 6**G). Our quantitative analysis using the Rosenthal & Henderson plot (**Fig. 6**H) revealed a significantly smaller B factor for VPP images compared to CEM images, suggesting that for CEM to achieve the same resolution as VPP it would necessitate significantly more particles, affirming a previously predicted quantitative benefit of using phase plate imaging for small proteins without symmetry [41].

Phase plate (PP) is a crucial microscope device that enhances phase contrast by altering the path of light or electron rays [44]. Since the advent of electron microscope PP [24], its potential has been identified for enabling visualization of many substances that were invisible because of limitations on imaging doses due to the radiation sensitivity. This capability of electron microscope PP is due to that it restores the low spatial frequency. As a result, electron microscope PP has found best application for cryo-EM imaging of biological macromolecules, in particular those with small size [25] or embedded in thick medium [30], or cells [45–46]. Our study harnessed capabilities of electron microscope PP across those aspects, and represents a unique case of employing two different thin-film PPs. While rigorous and fair comparisons of the performance of ZPP and VPP are challenging due to their installation being on distinct microscope platforms, it is beneficial to briefly outline the strengths and weaknesses of them (please also see [47]). The carbon film ZPP utilizes a pinhole to let the un-scattered beam through, with the scattered beam passing through the surrounding film to maximize the phase contrast through a 90-degree phase shift between the two beams. In contrast, VPP, made of a continuous film, presents a hole-free solution [31] where the phase shift between the un-scattered and scattered beams is produced by an in situ generated Volta field [32], which in turns retards the un-scattered beam. Notably, the Volta phase contrast gradually increases as the Volta field builds up. As for an ideal ZPP electron microscope, when it is operated with objective lens set at in-focus conditions together with minimal spherical aberration (Cs), it can provide in-phase structural information up to the regime of atomic resolution, abolishing the need of CTF correction. However, in our case we employed a customer designed ZPP cryo-EM with a fairly large Cs (4.2 mm) and operated it at close to focus condition to target medium-resolution information (∼10 Å) [26]. Due to practical challenges with use of this film ZPP, including poor yield of good pinholes (10-20%), difficulties in manual pinhole alignment with keeping it on axis, and pinhole charging, this ZPP cryo-EM has found limitations in its application in scaling up images required for fruitful single-particle analysis. By contrast, VPP did not face those issues, and its applications further benefit from cryo-EM automation and CTF correction [34]. As a result, our studies with VPP not only succeeded in replicating most advantages with ZPP in terms of identifying IgG in DDM and detecting small Fab particles in noisy cryo-EM images, but also in efficiently scaling up images to yield 3D structures of IgG and Fab with sub-nanometer resolutions. These advancements with VPP seemed to eclipse the need for ZPP in our pilot experiments. On reflection, the detour to ZPP on a multi-purposed instrument provided valuable preliminary data, supporting the rationale for gaining approval to use VPPs on automated cryo-EMs in our core facility. This approval might have been denied based on the mass limit empirically established by Herzik Jr et al [21]. In any case, our study suggests those mass limits can be further pushed with PP contrast where effective utilization of superior ZPP contrast can be obtained from a new type of ZPP [48] free from those issues with carbon film.

In summary, our cryo-EM study has harnessed phase contrast microscope techniques to address challenges posed by embedding-medium or protein size. This approach has yielded the first cryo-EM structure of a therapeutic IgG in isolation, achieving sub-nanometer resolution. Our findings provide previously unknown structural insights into the diverse conformations of therapeutic IgG in solution, laying a foundation for future measuring conformation landscape of IgG drug that bears pharmaceutical interests. Furthermore, our investigation into Fab has underscored the pivotal role of phase plates in cryo-EM structural analysis, especially for sub-60 kDa proteins that lack symmetry. The demonstrated imaging and structural analysis workflow is not only useful for therapeutic IgG research but is also broadly applicable to other biological molecules with similar challenging features.

## Materials and Methods

### Sample preparation for Dynamical Light Scattering (DLS) Measurement

Rituximab purchased from vendor (10mg/ml in PBS) was diluted to 2 mg/ml with PBS or deionized water. When n-Dodecyl-beta-Maltoside (DDM) was included for comparison, it was used at 0.008% (w/v), which was below its critical micelle concentration (CMC) of 0.0087%. DLS experiments were performed with DynaPro NanoStar (Wyatt Technology, Santa Barbara, CA, USA)

### Sample Preparation for Cryo-EM

Cryo-EM sample preparation was carried out immediately after the protein was diluted. For the Rituximab sample, vendor supplied protein (10 mg/ml in PBS) was first diluted to 0.5 mg/ml or 2 mg/ml with deionized water. Further addition of DDM was added to the water-diluted Rituximab at 0.008% (w/v), which is below its critical micelle concentration (CMC) of 0.0087%. 3 μl diluted protein sample was put on freshly glow-discharged Quantifoil R1.2/1.3 Cu grid (Quantifoil Micro Tools GmbH, Großlöbichau, Germany). With Vitrobot (Model Mark IV, FEI Company, Hillsboro, OR, USA), the protein-loaded grid was mounted on the plunger in the chamber kept at 4 °C and 98% humidity. The grid was hanged for 10 sec followed by blotting for 4.5 sec with blotting force set to 0, and then quickly plunged into liquid ethane for freezing. The made cryo-specimen was either immediately used for imaging or stored in liquid nitrogen for later use.

### Fab cleavage

The Fab fragment of Rituximab was prepared by using the PierceTM Fab preparation kit (Thermo Fisher Scientific Inc., Waltham, MA) according to the instructions. Briefly, the buffer of Rituximab (0.5 mg) was first exchanged with the papain digestion buffer and adjusted to a final volume of 0.5 ml. The prepared Rituximab was added to a spin column tube containing the resin with immobilized papain and incubated at 37° C for 3 h. The spin column tube was then centrifuged at 5000 x g for 1 min to collect the digested Rituximab, and the buffer was exchanged again to PBS. The Fab digested from Rituximab was separated from Fc using Protein A plus spin column, and collected in the flow-through. Protein concentration was measured to be 0.1 mg/ml by the BCA protein assay kit (Thermo Fisher Scientific Inc., Waltham, MA). To make cryo-specimen of Fab fragment, the protein was diluted from 0.1 mg/ml to 0.05 mg/ml with deionized water and loaded onto a freshly glow-discharged Quantifoil R1.2/1.3 Cu grid (Quantifoil Micro Tools GmbH, Großlöbichau, Germany) followed by plunge-freezing using Vitrobot as above-mentioned.

### Conventional cryo-EM

Conventional imaging of Rituximab diluted with deionized water to 1 mg/ml was performed with a 200 kV automated cryo-EM with a thermal field emission electron source (Cryo-ARM, JEOL Ltd., Tokyo, Japan) at Osaka University. Conventional imaging of diluted Rituximab (0.5 mg/ml/) with addition of DDM (0.008%) was performed with a 200 kV multi-purpose cryo-EM with a thermal field emission electron source (FS2200, JEOL Ltd., Tokyo, Japan) at Okazaki. Frozen grids transferred to an automated 200 kV cryo-EM (Cryo-ARM, JEOL Ltd., Tokyo, Japan) were via a grid cassette (JEOL Ltd., Tokyo, Japan), while those to JEM-FS2200 200 kV (JEOL Ltd., Tokyo, Japan) were via a Model 626 holder (Gatan Ltd., Pleasanton, CA, USA). The imaging on JEM-FS2200 was recorded manually on DE-20 direct electron detector (DDD Ltd, USA) whereas on cryo-ARM was on K2 direct electron detector (Gatan Ltd, Pleasanton, CA, USA) using JADAS software (JEOL Ltd., Tokyo, Japan). The nominal magnification in both cases was set to 40,000×. Each image movie recorded on DE-20 by FS2200 consisted of 75 movie frames in 3 seconds with the pixel size corresponded to 1.4 Å, and the accumulated electron doses on the specimen were approximately 40 e^−^/Å^2^ (**Table 1**). Each image movie recorded on K2 by Cryo-ARM contained 40 frames in 8 seconds with the pixel size corresponded to 1.09 Å, and the accumulated electron doses on the specimen were 50 e^−^/Å^2^. Movie frames were motion-corrected using MotionCorr2 and summed [50].

### Zernike phase plate multi-purpose cryo-EM

Frozen grids were transferred via a side-entry cryo-holder (Model 626, Gatan Ltd, Pleasanton, CA, USA) and imaged with a 200 kV multi-purpose cryo-EM with an omega-type energy filter and field emission electron source (JEM-2200FS, JEOL Ltd., Tokyo, Japan). The images were recorded on a DE-20 direct detector (Direct Electron LP, San Diego, CA, USA) with nominal magnification of 40,000×. Arrays of Zernike phase plate (ZPP) pinholes [49] were fabricated in NIPS laboratory, and installed in the back focal plane of the objective lens (Cs 4.2 mm) one day before the use for imaging experiments. Positioning a ZPP pinhole was done by moving it on the X-Y plane manually to the optical axis as the back focal plane was being viewed, followed by Z adjustment to achieve the on-plane condition [49] by tweaking the C2 lens to make the pinhole image slightly larger than that of a Quantifoil hole followed by insertion of the energy filter and image recording. Each image contained 72 movie frames in 3 seconds was collected in-focus with pixel size corresponding to 1.4 Å on the specimen. The accumulated electron doses on the specimen for each image were approximately 40 e^−^/Å^2^ (**Table 1 & 2**). Normally, 10 to 30 images could be obtained from a ZPP position before it was no longer usable due to plate charging issues. Movie frames were motion-corrected using MotionCorr2 and summed [50].

### HFPP multi-purpose cryo-EM

Hole-free phase plate (HFPP) imaging was manually performed at JEOL factory. With a frozen grid transferred to a JEM-F200 multi-purpose cryo-EM with cold field emission source (JEOL Ltd., Tokyo, Japan) via a side-entry cryo-holder (Model 626, Gatan Ltd, Pleasanton, CA, USA). To develop a Volta field on HFPP with JEM-F200 usually took 3-5 minutes while the on-plane condition was found by Ronchigram method. The images were recorded in a near in-focus condition on a K2 direct detector (Gatan Ltd, Pleasanton, CA, USA) with nominal magnification of 40,000× (**Table 1**). Each image contained 40 movie frames was collected in 8 seconds with a pixel size corresponding to 1.09 Å and accumulated electron dose a on the specimen were approximately 50 e^−^/Å^2^. Movie frames were motion-corrected using MotionCor2 and summed [50].

### Volta PP automated cryo-EM imaging

To facilitate phase plate cryo-EM with automated data collection of full-length Rituximab, over-night imaging session was performed using 200 kV Talos Artica cryo-EM (Thermo Fisher Scientific, USA) with Volta phase plate. The images were recorded on direct electron camera (Falcon III, Thermo Fisher Scientific, USA). To develop a Volta field on carbon film phase plate, 15 second of pre-charging process was applied when a large C2 aperture (150 micron) was used while the on-plane condition was found by Ronchigram method. This pre-charging had primed the initial phase shift to 30 degree before data collection started. Each Volta phase plate (VPP) position was used for collecting only 20 images to control the phase shift from being further developed beyond 90 degree to prevent the associated plate charging issues. Post-imaging phase shift analysis showed that the phase shift could occasionally go beyond 90 degree to reach 110 degree. Data acquisition was automatically performed using FEI EPU software at a nominal magnification of 120,000×, which corresponded to a pixel size of 0.855 Å per pixel (**Table 1**). Each 2.5-second movie consisting of 50 frames was collected using linear mode to yield 50 e^−^/Å^2^ on the specimen. A total of 606 images were obtained in one session (**Table 1**).

For automated cryo-EM imaging of Fab fragment, we employed Titan Krios with K3 direct electron camera (Gatan Ltd, Pleasanton, CA, USA). Imaging session was performed with Volta phase plate. After 30 seconds pre-charging, the initial phase shift was primed at 40 degree prior to movie data collection. Each VPP position was used to collect 30 images. Automated Data acquisition was performed using FEI EPU software where the nominal magnification was set to 105,000x using super-resolution mode, corresponding to pixel size of 0.415 Å (**Table 2**). Each image consisting of 50 movie frames was collected within 1.8 seconds to yield accumulated dose of 50 e^−^/Å^2^ on the specimen. A total of 956 images were obtained (**Table 2**). For comparative conventional cryo-EM imaging of Fab, the same magnification and dose parameters were used where the defocus was set in the range of 1.6 to 3 microns, and a total of 10,166 movies were collected (**Table 2**).

### Cryo-EM image analysis

For the case of full-length Rituximab, stacks of 606 Talos Acrtica movies were sequentially processed using MotionCorr2 [50] for motion correction, and Gctf [51] was used for CTF estimation. Single particle analysis was performed with cryoSPARC v4.1 [36]. Particles were first auto-picked from all micrographs by Blob Picker with particle diameter set to be 150Å. Subsequently, one round of 2D classification was performed. A few clear 2D class averages were obtained, and used as templates for particle re-picking from all micrographs using Template Picker. A total of 173,336 particles were picked by this re-picking process, and subjected to several rounds of reference-free 2D classification, by which a final set of 31,111 particles was obtained. Those 31,111 particles were employed for building a 3D ab-initio model. A 3D model with three-arm structure was used as a reference for performing “3D Homogeneous Refinement” to improve resolution of this three-arm structure. To improve the resolution for the Fab domain, focused refinement was applied to a Fab arm by making it. Subsequently, 3D variability analysis (3DVA) [39] was applied to the set of 31,111 particles to test whether or not multiple conformations could be disentangled. 3D structures were then displayed and presented using Chimera [52].

For the case of imaging Fab fragment, cryo-EM images were obtained from Titan Krios with VPP with defocusing. A stack of 956 movies was obtained with VPP and defocusing [34], pre-processed with MotionCorr2 and Gctf for CTF correction followed by auto-picking using RELION (3.1.2). 288,931 particles were picked with a tight threshold to reduce the chances of false picking of non-particle artifacts associated with background noise; 171,288 particles were obtained after several rounds of 2D Classification. Subsequently, many rounds of 3D ab-initio model generation were attempted with sub-sampling the set of 171,288 particles using cryoSPARC (v4.1). A reliable subset of 72,035 particles was identified based on their ability to give rise to a low-resolution 3D ab-initio model with the characteristic two-lobe feature. This subset of particles together with the corresponding 3D ab-initio model as a reference model was employed for refinement using “Non-uniform Refinement” in cryoSPARC v4.1. 3D structures were displayed and presented using Chimera [52].

## Additional Information

### Conflict of Interest

The authors declare no competing interests.

## Acknowledgements

This project was supported by Academia Sinica SUMMIT Project [AS-SUMMIT-107], [AS-SUMMIT-108], [AS-SUMMIT-109], and Academia Sinica Seed Grant [AS-GCS-112-M03] to W.-H.C.; I-P.T. was supported by Academia Sinica Seed Grant [AS-GCS-108-08], and an Academia Sinica Investigator Award [AS-IA-110-M05]. The imaging of cryo-EM at Academia Sinica was helped by Dr. Kondo Yuan-Chih Chang while its routine operation was supported by [AS-CFII-108-110]. The authors thank Dr. Chris Shu-Chuan Jao and Mr. Xin Jie Huang in the Biophysics Core Facility (BCF) Academia Sinica, for their technical assistance and analytical advice on using DynaPro NanoStar for DLS experiments. The BCF is funded by Academia Sinica Core Facility and Innovative Instrument Project (AS-CFII-111-201). The operation of Cryo-EM of Okazaki National Institute for Physiological Sciences (NIPS) was supported by NIPS joint research grants. The authors thank JEOL factory for using F-200 field emission multi-purpose microscope, and are grateful for assistance of Dr. Chia-Ling Chen in Che Ma lab, Genomic Research Center of Academia Sinica, on initial preparation of antibody for imaging tests at Osaka University in May 2017 that helped launch this project.

## Data Availability

Raw data of cryo-EM images are available from the corresponding author (W.-H.C.) upon reasonable request. The cryo-EM structure of Rituximab IgG has been deposited on PDB databank with accession number EMD-38153; and the VPP Fab and CEM Fab structures are deposited with EMD-38154, and EMD-38155, respectively.

## Author Contributions

W.-H. Chang and C.-H. Wong conceived this project. C.-H.Y. and T.-L.H. prepared Rituximab sample and performed enzymatic cleavage for Fab fragmentation. C.-H. Wang, T.K., and K.N. performed imaging on Cryo-ARM (JEOL, Japan) at Osaka University. C.-H. Wang, S.-H.H., and K.H. performed HFPP imaging on F-200 (JEOL, Japan) at JEOL factory. H.-H.L., S.-H.H., C.S. and K.M. performed Zernike cryo-EM imaging on FS-2200 (JEOL, Japan) at Okazaki. H.-H.L. and C.-H. Wang. performed Volta cryo-EM imaging on Talos Arctica (Thermo Fisher Scientific, U.S.A.) at Academia Sinica. H.-H.L. and C.-H. Wang. performed conventional cryo-EM imaging on Titan Krios (Thermo Fisher Scientific, U.S.A.) at Academia Sinica. H.-H.L. and Y.-M.W. performed Volta cryo-EM imaging on Titan Krios (Thermo Fisher Scientific, U.S.A) at Academia Sinica. S.-H.H. and S.S.-Y.L. performed membrane protein expression and detergent addition experiments. H.-H.L., I-P.T. and W.-H.C. analyzed the data. H.-H. L. and W.-H.C. wrote the manuscript.

## Notes

### Competing Interest Statement

The authors have declared no competing interest.

