## SUPPLEMENTARY INFORMATION for "Use of phase plate cryo-EM reveals conformation diversity of therapeutic IgG with 50 kDa Fab fragment resolved below 6 Å"

### for

Present Addresses:

**This file contains 6 supplementary figures**

**Supplementary Figure 1**

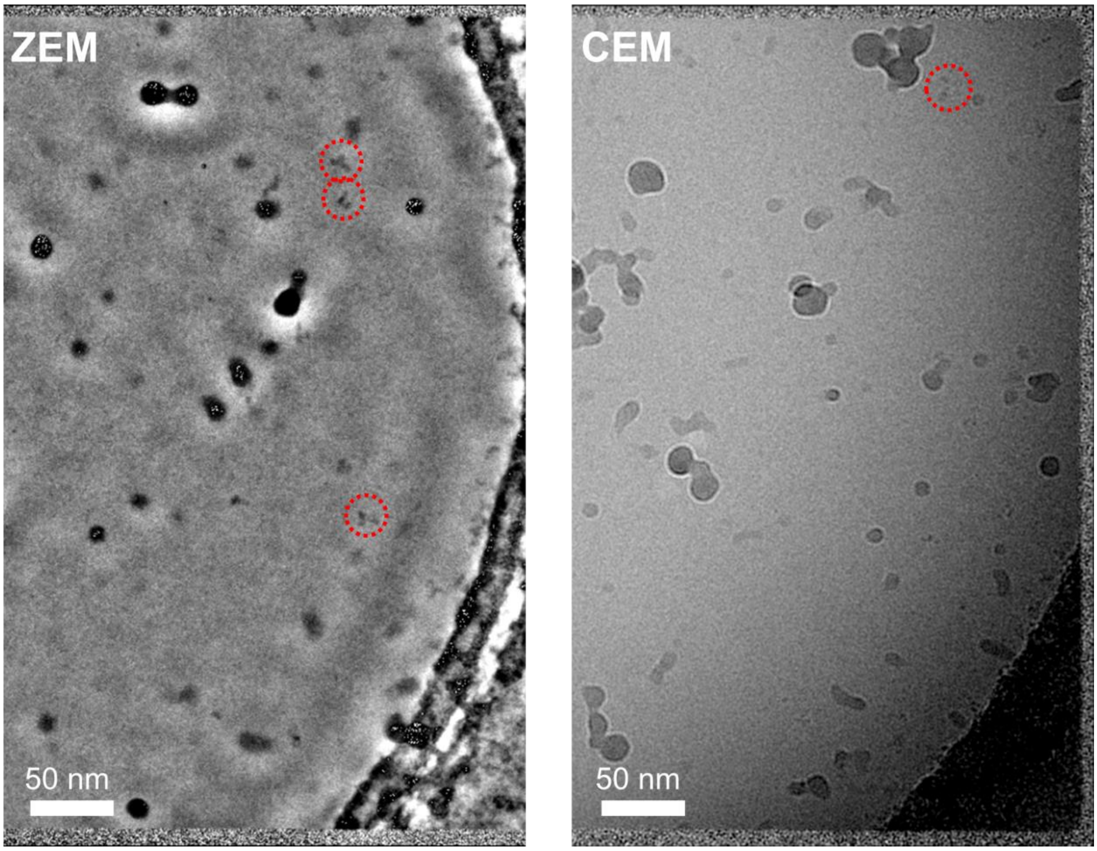


**Figure S1. DDM-dispersed Rituximab IgG under phase plate and conventional cryo-EM.** The left is a Zernike phase plate cryo-EM (ZEM) image of DDM-solubilized Rituximab (0.5 mg/ml with 0.008% DDM) recorded by a direct electron camera (DE-20, DDD Ltd, USA) on a FS-2200 multi-purpose 200 kV cryo-EM (JEOL Ltd, Japan) where several three-blob particles are identified and denoted by red circles. Shown in the right is a conventional cryo-EM image of the same DDM-solubilized Rituximab specimen recorded by the same camera on the same microscope where a candidate Rituximab antibody is highlighted with a red circle.

**Supplementary Figure 2**

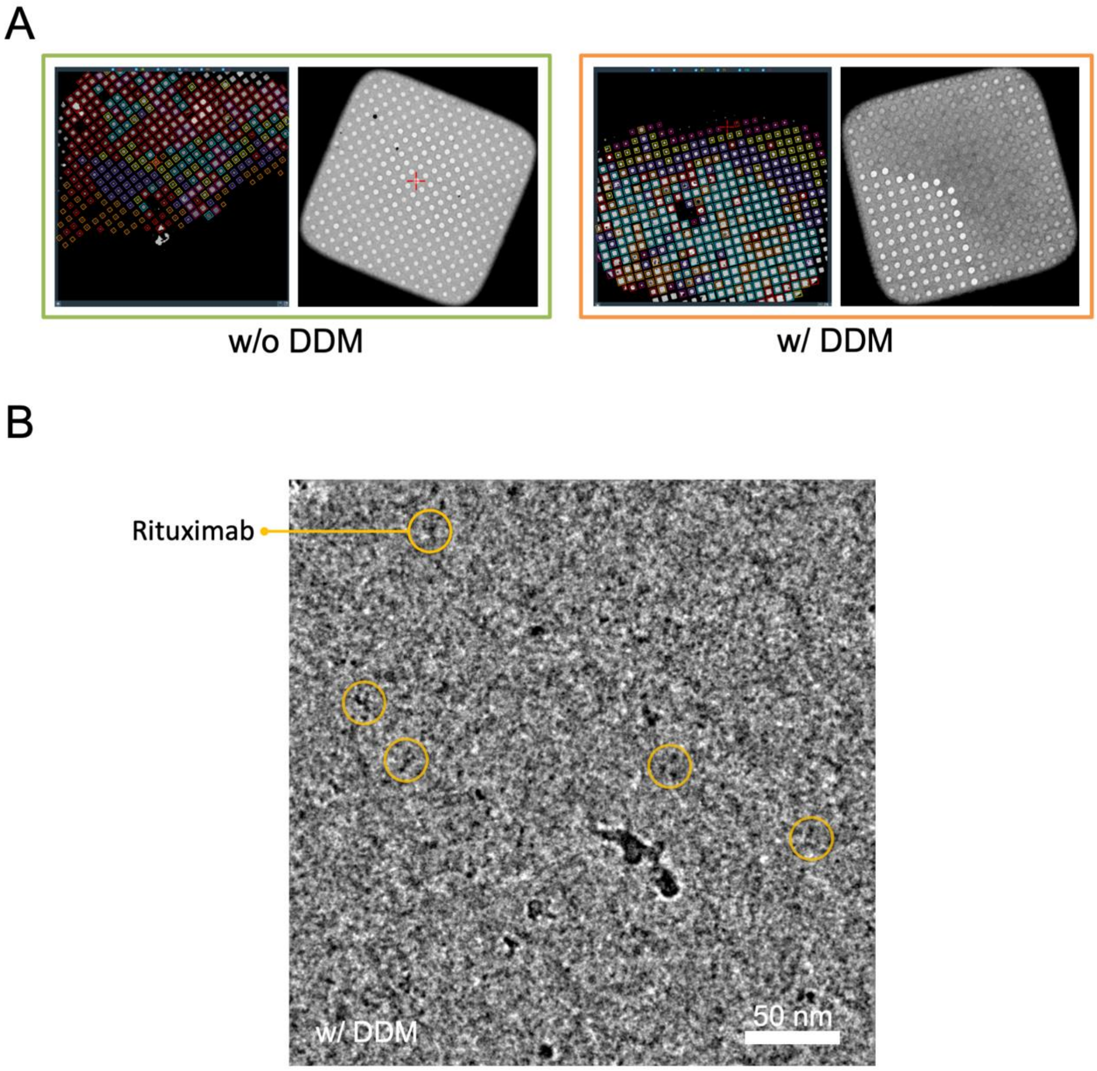


**Figure S2. Low magnification atlas of entire grid with a representative grid square of two cryo-EM specimens, one without and the other with DDM** (0.008%)**. (A)** Without DDM, the gray scale that reflects the ice thickness in a grid square shows the ice thickness in all holes is uniformly distributed and on the thinner side compared to that in (B). **(B)** With DDM (0.008%), the gray scales in a grid square become diversified where ~70% of holes are apparently thicker. **(C)** A high magnification micrograph of specimen of Rituximab with DDM (0.008%) was recorded by conventional cryo-EM technique with -3µm defocus value. A Rituximab molecule is circled where the background is very noisy. All images were recorded on Talos Arctica (Thermo Fisher, USA).

**Supplementary Figure 3**

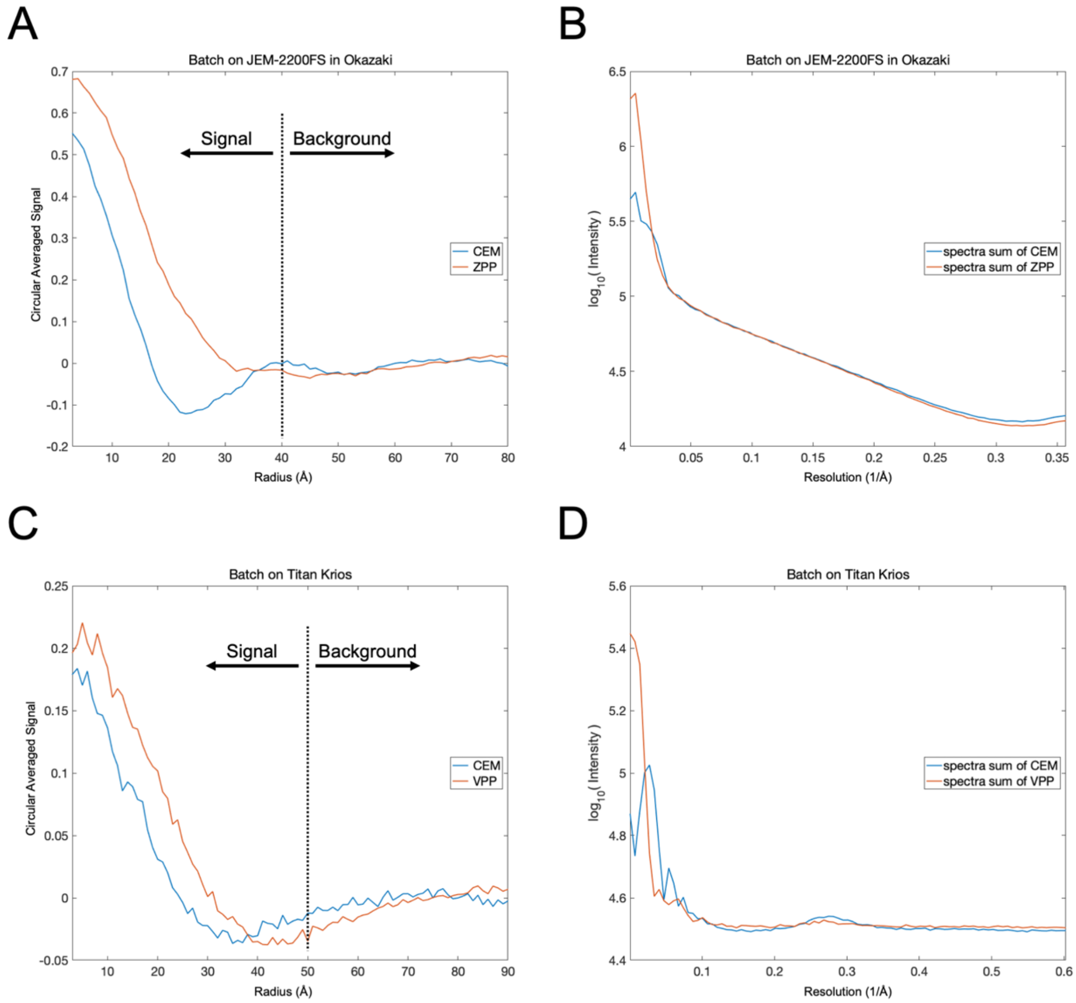


**Figure S3. Contrast enhancement by phase plates characterized by signal to background in cryo-EM image and spatial frequency in Fourier transform. (A)** Signal to background of ZPP Fab image (in red) and CEM Fab image (in blue). The particle profile is obtained from rotational averaging and plotted against particle radius. The images were taken by a direct electron camera (DE-20, DDD, USA) on FS-2200 200 kV multi-purpose cryo-EM (JEOL, Japan). **(B)** Comparison of ZPP and CEM Fourier transforms. ZPP transform has 5 times higher amplitude than CEM with 3 micron defocus) at very low spatial frequency. **(C)** Signal to background of VPP image (in red) and CEM image (in blue) as plotted against particle radius. The images were taken by a direct electron camera (K3, Gatan, USA) on Titan Krios 300 kV cryo-EM (Thermo Fisher, USA). **(D)** Comparison of VPP and CEM Fourier transforms shows at very low spatial frequency regime, the VPP has 3 times higher amplitude than CEM (3 micron defocus).

**Supplementary Figure 4**

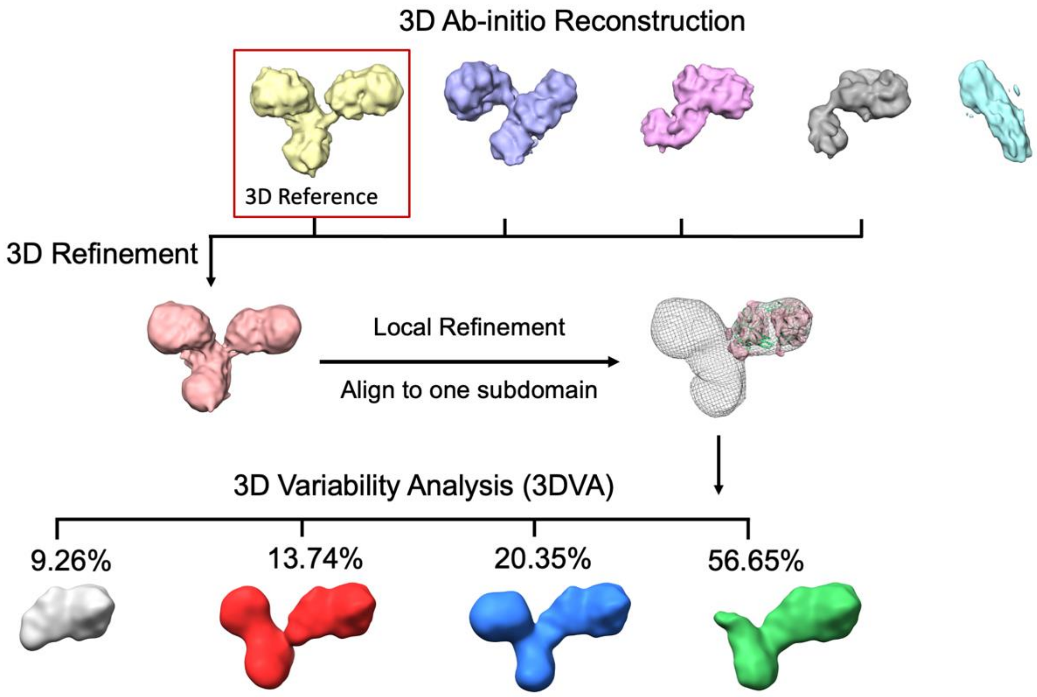


**Figure S4.** A workflow of 3D reconstruction for Rituximab IgG from ab initio using VPP cryo-EM images using cryoSPARC. Each toss had resulted in a different initial model, where the frequency of getting the three-arm structure is ~30%. 3D refinement were carried out in two stages, global refinement that optimized the alignment parameters of each IgG particle image, and local refinement (focused refinement) that focus on the alignment of the Fab domain by masking it in each IgG image. Further 3DVA analysis, also using cryoSparc to resolve heterogeneous conformations. A total of four meaningful conformations are obtained, two are with three arms (red and blue), and one with two arms (green), and one with one arm (grey).

**Supplementary Figure 5**

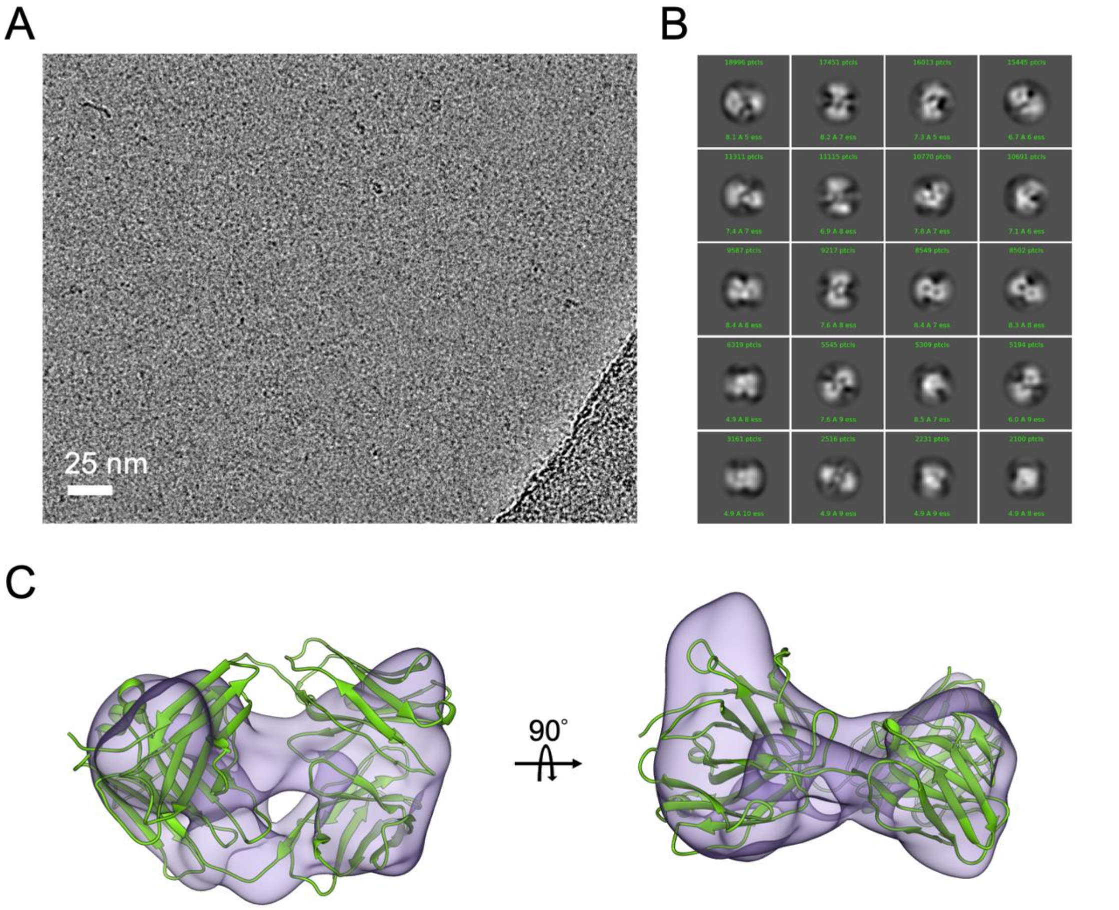


**Figure S5. Conventional cryo-EM imaging of Rituximab Fab fragment. (A)** A defocused CEM image shows Fab particles with highly noisy background. **(B)** Representative 2D class averages. **(C)** 3D reconstruction of Fab fragment with docking of X-ray model (PDB:4KAQ).
